# *INPP5D* expression is associated with risk for Alzheimer’s disease and induced by plaque-associated microglia

**DOI:** 10.1101/2020.08.31.276444

**Authors:** Andy P. Tsai, Peter Bor-Chian Lin, Chuanpeng Dong, Miguel Moutinho, Brad T. Casali, Yunlong Liu, Bruce T. Lamb, Gary E. Landreth, Adrian L. Oblak, Kwangsik Nho

**Affiliations:** Stark Neurosciences Research Institute, IUSM, Indianapolis, IN, USA. (A.P.T); (P.B.L); (M.M); (G.E.L); (B.T.L); (A.L.O); Department of Medical and Molecular Genetics, Center for Computational Biology and Bioinformatics, IUSM, Indianapolis, IN, USA. (C.D); (Y.L); Department of Neurosciences, Case Western Reserve University, School of Medicine, Cleveland, OH, USA. (B.T.C); Department of Medical and Molecular Genetics, IUSM, Indianapolis, IN, USA. (B.T.L); Department of Anatomy and Cell Biology, IUSM, Indianapolis, IN, USA. (G.E.L); Department of Radiology & Imaging Sciences, IUSM, Indianapolis, IN, USA. (A.L.O); (K.N)

**Keywords:** Alzheimer’s disease (AD), Microglia, INPP5D, AD risk, Plaque

## Abstract

**Background:** Alzheimer’s disease (AD) is a progressive neurodegenerative disorder characterized by cognitive decline, robust microgliosis, neuroinflammation, and neuronal loss. Genome-wide association studies recently highlighted a prominent role for microglia in late-onset AD (LOAD). Specifically, inositol polyphosphate-5-phosphatase (*INPP5D*), also known as SHIP1, is selectively expressed in brain microglia and has been reported to be associated with LOAD. Although *INPP5D* is likely a crucial player in AD pathophysiology, its role in disease onset and progression remains unclear.

**Methods:** We performed differential gene expression analysis to investigate *INPP5D* expression in LOAD and its association with plaque density and microglial markers using transcriptomic (RNA-Seq) data from the Accelerating Medicines Partnership for Alzheimer’s Disease (AMP-AD) cohort. We also performed quantitative real-time PCR, immunoblotting, and immunofluorescence assays to assess INPP5D expression in the 5xFAD amyloid mouse model.

**Results:** Differential gene expression analysis found that *INPP5D* expression was upregulated in LOAD and positively correlated with amyloid plaque density. In addition, in 5xFAD mice, *Inpp5d* expression increased as the disease progressed, and selectively in plaque-associated microglia. Increased *Inpp5d* expression levels in 5xFAD mice were abolished entirely by depleting microglia with the colony-stimulating factor receptor-1 antagonist PLX5622.

**Conclusions:** Our findings show that *INPP5D* expression increases as AD progresses, predominantly in plaque-associated microglia. Importantly, we provide the first evidence that increased *INPP5D* expression might be a risk factor in AD, highlighting *INPP5D* as a potential therapeutic target. Moreover, we have shown that the 5xFAD mouse model is appropriate for studying *INPP5D* in AD.

## Background

Alzheimer’s disease (AD) is the most common cause of dementia, with pathogenesis arising from perturbed β-amyloid (Aβ) homeostasis in the brain [1]. The mechanisms underlying the development of the most common form of AD, late-onset AD (LOAD), are still unknown. Microglia, the primary immune cells in the brain play a crucial role in AD pathogenesis [2]. Recent large-scale genome-wide association studies (GWAS) reported that many genetic loci associated with LOAD risk are related to inflammatory pathways, suggesting that microglia are involved in modulating AD pathogenesis [3, 4]. Among the microglia-related genetic factors in LOAD, a common variant in *INPP5D* (phosphatidylinositol 3,4,5-trisphosphate 5-phosphatase 1), rs35349669, confers an increase in LOAD risk (OR=1.08) [4, 5]. Conversely, the intronic *INPP5D* variant rs61068452 is associated with a reduced CSF t-tau/Aβ1-42 ratio, plays a protective role in LOAD (p=1.48E-07) [6]. *INPP5D* encodes inositol polyphosphate-5-phosphatase which participates in regulation of microglial gene expression [7]. Specifically, *INPP5D* inhibits signal transduction initiated by activation of immune cell surface receptors, including Triggering receptor expressed on myeloid cells 2 (TREM2), Fc gamma receptor (Fc*γ*R) and Dectin-1 [8]. The conversion of PI(3,4,5)P3 to PI(3,4)P2 is catalyzed by INPP5D following its translocation from the cytosol to the cytoplasmic membrane. The loss of PI(3,4,5)P3 prevents the activation of the immune cell surface receptors [9]. Interestingly, genetic variants of TREM2, Fc*γ*R, and Dectin-1 are also associated with increased AD risk [10–12] and are potentially involved in regulating INPP5D activity. Inhibiting INPP5D promotes microglial proliferation, phagocytosis, and increases lysosomal compartment size [13]. Although INPP5D has been shown to play an important role in microglial function, its role in AD remains unclear.

Here, we report that *INPP5D* is upregulated in LOAD, and elevated *INPP5D* expression levels are associated with microglial markers and amyloid plaque density. Furthermore, in the 5xFAD mouse model, we found a disease-progression-dependent increase in *INPP5D* expression in plaque-associated microglia. Our results suggest that *INPP5D* plays a role in microglia phenotypes in AD and is a potential target for microglia-focused AD therapies.

## Methods

### Human participants and RNA-Seq

RNA-Seq data were obtained from the AMP-AD Consortium, including participants of the Mayo Clinic Brain Bank cohort, the Mount Sinai Medical Center Brain Bank (MSBB) cohort, and the Religious Orders Study and Memory and Aging Project (ROSMAP) cohort.

In the Mayo Clinic RNA-Seq dataset [14], the RNA-Seq-based whole transcriptome data were generated from human samples of 151 temporal cortices (TCX) (71 cognitively normal older adult controls (CN) and 80 LOAD) and 151 cerebella (CER) (72 CN and 79 LOAD). LOAD participants met the neuropathological criteria for AD (Braak score ≥4.0), and cognitively normal participants had no neurodegenerative diagnosis (Braak score ≤ 3.0).

In the MSBB dataset [15], data were generated from human samples from CN, mild cognitive impairment (MCI), and LOAD participants’ parahippocampal gyrus (PHG) and inferior frontal gyrus (IFG), superior temporal gyrus (STG) and frontal pole (FP). The clinical dementia rating scale (CDR) was used to assess dementia and cognitive status [16]. LOAD patients had a CDR ≥0.5, while MCI and CN participants had a CDR of 0.5 and 0, respectively. CN participants had no significant memory concerns. This study included 108 participants (16 CN, 14 MCI, and 78 LOAD) for PHG, 137 participants (21 CN, 18 MCI, and 98 LOAD) for STG, 136 participants (18 CN, 16 MCI, and 102 LOAD) for IFG, and 153 participants (22 CN, 20 MCI, and 111 LOAD) for FP.

In the ROSMAP dataset [17], RNA-Seq data were generated from the dorsolateral prefrontal cortices of 241 participants (86 CN and 155 LOAD).

### Animal models

Wild-type (WT) and 5xFAD mice were maintained on the C57BL/6J background (JAX MMRRC Stock# 034848) for IHC and qPCR studies. Two-, four-, six-, eight-, and twelve-month-old mice were used. In the PLX5622 study, we used WT and 5xFAD mice maintained on the mixed C57BL/6J and SJL background [B6SJL-Tg (APPSwFlLon, PSEN1*M146L*L286V) 6799Vas, Stock #34840-JAX]) (**Fig. 3e and 3f**). The 5XFAD transgenic mice overexpress five FAD mutations: the APP (695) transgene contains the Swedish (K670N, M671L), Florida (I716V), and London (V717I) mutations and the PSEN1 transgene contains the M146L and L286V FAD mutations. Up to five mice were housed per cage with SaniChip bedding and LabDiet® 5K52/5K67 (6% fat) feed. The colony room was kept on a 12:12 hr. light/dark schedule with the lights on from 7:00 am to 7:00 pm daily. They were bred and housed in specific-pathogen-free conditions. Both male and female mice were used.

### PLX5622 animal treatment

At four months of age, either normal rodent diet or PLX5622-containing chow was administered to 5XFAD mice for 28 days. An additional cohort of four-month-old mice was treated with PLX5622 or control diet for 28 days, then discontinued from PLX5622 feed and fed a normal rodent diet for an additional 28 days. At six months of age, this cohort of mice was euthanized. Plexxikon Inc. provided PLX5622 formulated in AIN-7 diet at 1200 mg/kg [18].

### Statistical analysis

In the human study, differential expression analysis was performed using *limma* software [19] to investigate the diagnosis group difference of *INPP5D* between CN, MCI, and LOAD. Age, sex, and *APOE ε4* carrier status were used as covariates. To investigate the association between *INPP5D* expression levels and amyloid plaque density or expression levels of microglia-specific markers *(AIF1* and *TMEM119*), we used a generalized linear regression model with *INPP5D* expression levels as a dependent variable and plaque density or microglia-specific markers along with age, sex, and *APOE ε4* carrier status as explanatory variables. The regression was performed with the “glm” function from the stats package in R (version 3.6.1).

In the mouse study, GraphPad Prism (Version 8.4.3) was used to perform the statistical analyses. Differential expression analysis of both gene and protein levels between WT and 5xFAD mice was performed using unpaired Student’s t-test. The statistical comparisons between mice with and without PLX5622 treatments were performed with one-way ANOVA followed by Tukey’s posthoc test. Graphs represent the mean and standard error of the mean.

### RNA extraction and quantitative real-time PCR

Mice were anesthetized with Avertin and perfused with ice-cold phosphate-buffered saline (PBS). The cortical and hippocampal regions from the hemisphere were micro-dissected and stored at −80°C. Frozen brain tissue was homogenized in buffer containing 20 mM Tris-HCl (pH=7.4), 250 mM sucrose, 0.5 mM EGTA, 0.5 mM EDTA, RNase-free water, and stored in an equal volume of RNA-Bee (Amsbio, CS-104B) at −80°C until RNA extraction. RNA was isolated by chloroform extraction and purified with the Purelink RNA Mini Kit (Life Technologies #12183020) with an on-column DNAse Purelink Lit (Life Technologies #12183025). 500 ng RNA was converted to cDNA with the High-Capacity RNA-to-cDNA Kit (Applied Biosystems #4388950), and qPCR was performed on a StepOne Plus Real-Time PCR system (Life Technologies). Relative gene expression was determined with the 2^-ΔΔCT^method and assessed relative to *Gapdh* (Mm99999915_g1). *Inpp5d* primer: Taqman Gene Expression Assay *(Inpp5d:* Mm00494987_m1 from the Life Technologies). Student’s *t*-test was performed for qPCR assays, comparing WT with 5xFAD animals.

### Immunofluorescence

Brains were fixed in 4% PFA overnight at 4°C. Following overnight fixation, brains were cryoprotected in 30% sucrose at 4°C and embedded. Brains were processed on a microtome as 30 μm free-floating sections. For immunostaining, at least three matched brain sections were used. Free-floating sections were washed and permeabilized in 0.1% Triton in PBS (PBST), followed by antigen retrieval using 1x Reveal Decloaker (Biocare Medical) at 85°C for 10 mins. Sections were blocked in 5% normal donkey serum in PBST for 1 hr. at room temperature (RT). The following primary antibodies were incubated in 5% normal donkey serum in PBST overnight at 4°C: IBA1 (Novus Biologicals #NB100-1028 in goat, 1:1000); 6E10 (BioLegend #803001 in mouse, 1:1000; AB_2564653); and SHIP1/INPP5D (Cell Signaling Technology (CST) #4C8, 1:500, Rabbit mAb provided by CST in collaboration with Dr. Richard W. Cho). Sections were washed and visualized using respective species-specific AlexaFluor fluorescent antibodies (diluted 1:1000 in 5% normal donkey serum in PBST for 1 hr. at RT). Sections were counterstained and mounted onto slides. For X-34 staining (Sigma, #SML1954), sections were dried at RT, rehydrated in PBST, and stained for ten mins at RT. Sections were then washed five times in double-distilled water and washed again in PBST for five mins. Images were acquired on a fluorescent microscope with similar exposure and gains across stains and animals. Images were merged using ImageJ (NIH).

### Immunoblotting

Tissue was extracted and processed as described above, then centrifuged. Protein concentration was measured with a BCA kit (Thermo Scientific). 50 μg of protein per sample was boiled in SDS-PAGE protein sample buffer for 10 mins at 95°C, loaded into 4-12% Bis-Tris gels (Life Technologies) and run at 100 V for 90 mins. The following primer antibodies were used: SHIP1/INPP5D (CST #4C8 1:500, Rabbit mAb) and GAPDH (Santa Cruz #sc-32233). Each sample was normalized to GAPDH, and the graphs represent the values normalized to the mean of the WT mice group at each time point.

## Results

### *INPP5D* expression levels are increased in LOAD

INPP5D is a member of the inositol polyphosphate-5-phosphatase (INPP5) family and possesses a set of core domains, including an N-terminal SH2 domain (amino acids 5-101), Pleckstrin homology-related (PH-R) domain (amino acids 292-401), lipid phosphatase region (amino acids 401-866) with C2 domain (amino acids 725-863), and C-terminal proline-rich region (amino acids 920-1148) with two SH3 domains (amino acids 969-974 and 1040-1051) (**Fig. 1a**). Differential expression analysis was performed using RNA-Seq data from seven brain regions from the AMP-AD cohort. Expression levels of *INPP5D* were increased in the temporal cortex (logFC=0.35, p=1.12E-02; **Fig. 1b**), parahippocampal gyrus (logFC=0.54, p=7.17E-03; **Fig. 1c**), and inferior frontal gyrus (logFC=0.44, p=2.33E-03; **Fig. 1d**) of LOAD patients with age and sex as covariates (**Table 1**). Interestingly, *INPP5D* expression was also found to be increased in the inferior frontal gyrus of LOAD patients compared with MCI subjects (logFC=0.45, p=6.76E-03; **Fig. 1d**). Results were similar when *APOE* ε4 carrier status was used as an additional covariate. *INPP5D* remained overexpressed in the temporal cortex (logFC=0.34, p=2.75E-02), parahippocampal gyrus (logFC=0.53, p=1.08E-02), and inferior frontal gyrus (logFC=0.42, p=4.35E-03) of LOAD patients. However, we did not find any differences between the diagnosis groups in the cerebellum, frontal pole, superior temporal gyrus, or dorsolateral prefrontal cortex (**Table 1**). To examine whether *INPP5D* was associated with microglia, we analyzed the association between *INPP5D* and microglia-specific marker genes *(AIF1* and *TMEM119). AIF1* and *TMEM119* were significantly associated with *INPP5D* expression levels in the parahippocampal gyrus *(AIF1:* β=0.4386, p=4.10E-07; *TMEM119:* β=0.7647, p=<2E-16), inferior frontal gyrus *(AIF1:* β=0.2862, p=6.36E-08; *TMEM119:* β=0.6109, p=<2E-16), frontal pole *(AIF1:* β=0.2179, p=4.53E-04; *TMEM119* β=0.5062, p=4.00E-15), and superior temporal gyrus *(AIF1:* β=0.3013, p=5.36E-07; *TMEM119:* β=0.6914, p=<2E-16) (**Table 2**).

**Fig 1.**
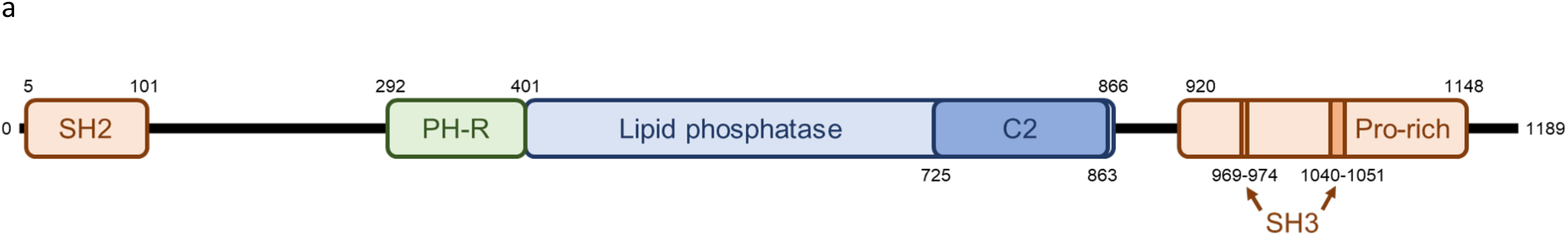
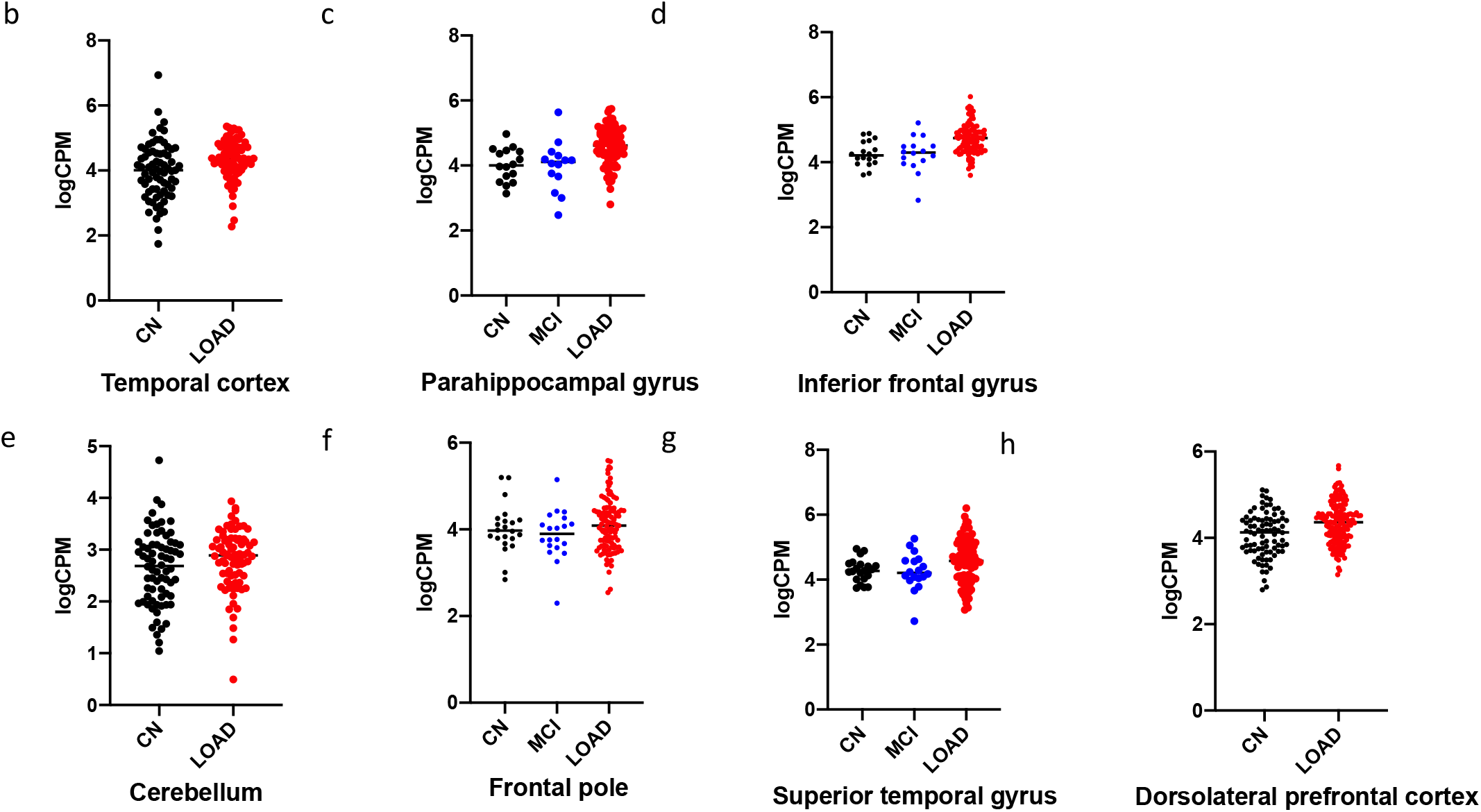
Relative quantification of *INPP5D* expression in the studied participants. (a) Domain architecture of *INPP5D* drawn to scale. Gene expression of *INPP5D* is showed as logCPM values in (b) Temporal cortex (TCX)-Mayo, (c) Parahippocampal gyrus (PHG)-MSBB, (d) Inferior frontal gyrus (IFG)-MSBB, (e) Cerebellum (CER)-Mayo, (f) Frontal pole (FP)-MSBB, (g) Superior temporal gyrus (STG)-MSBB, (h) Dorsolateral prefrontal cortex (DLPFC)-ROSMAP. SH2 *Src Homology 2 domain*, SH3 *SRC Homology 3 domain*, C2 *C2 domain* CN *cognitively normal*, MCI *mild cognitive impairment*, LOAD *Late-Onset Alzheimer’s disease*

**Table 1.** *INPP5D* expression levels were increased in LOAD. Table 1 shows the p-values for the gene expression analyses performed with *limma* using RNA-Seq data from the AMP-AD Consortium. TCX *temporal cortex*, PHG *parahippocampal gyrus*, STG *superior temporal gyrus*, IFG *inferior frontal gyrus*, FP *frontal pole*, CER *cerebellum*, DLPFC *dorsolateral prefrontal cortex*, CN *cognitively normal*, AD *Alzheimer’s disease*, MCI *mild cognitive impairment*, logFC *log fold-change*

| Brain Regions | Temporal Cortex | Parahippocampal Gyrus |  |  | Inferior Frontal Gyrus |  |  |
| --- | --- | --- | --- | --- | --- | --- | --- |
|  |  | Covariate: Age and Sex |  |  |  |  |  |
| Contrast | CN vs. LOAD | CN vs. MCI | MCI vs. LOAD | CN vs. LOAD | CN vs. MCI | MCI vs. LOAD | CN vs. LOAD |
| logFC | 0.34916854 | 0.012305567 | 0.565288103 | 0.536580956 | -0.009944963 | 0.44545717 | 0.43695423 |
| p-value | **1.12E-02** | 9.99E-01 | 7.56E-02 | **7.17E-03** | 1.00E+00 | **6.76E-03** | **2.33E-03** |
|  |  | Covariate: Age, Sex, and APOE ε4 status |  |  |  |  |  |
| Contrast | CN vs. LOAD | CN vs. MCI | MCI vs. LOAD | CN vs. LOAD | CN vs. MCI | MCI vs. LOAD | CN vs. LOAD |
| logFC | 0.34072691 | -0.034336626 | 0.57213088 | 0.52809965 | -0.0546836 | 0.46476734 | 0.42180431 |
| p-value | **2.75E-02** | 1.00E+00 | 1.00E-01 | **1.08E-02** | 1.00E+00 | **8.00E-03** | **4.35E-03** |

| Brain Regions | Cerebellum | Frontal Pole |  |  | Superior Temporal Gyrus |  |  | Dorsolateral Prefrontal Cortex |
| --- | --- | --- | --- | --- | --- | --- | --- | --- |
|  |  | Covariate: Age and Sex |  |  |  |  |  |  |
| Contrast | CN vs. LOAD | CN vs. MCI | MCI vs. LOAD | CN vs. LOAD | CN vs. MCI | MCI vs. LOAD | CN vs. LOAD | CN vs. LOAD |
| logFC | 0.15322052 | -0.1498388 | 0.20026554 | 0.09349678 | -0.0115078 | 0.31887371 | 0.28945778 | 0.1761858 |
| p-value | 2.15E-01 | 9.85E-01 | 4.80E-01 | 8.65E-01 | 1.00E+00 | 2.09E-01 | 1.78E-01 | 5.76E-02 |
|  |  | Covariate: Age, Sex, and APOE ε4 status |  |  |  |  |  |  |
| Contrast | CN vs. LOAD | CN vs. MCI | MCI vs. LOAD | CN vs. LOAD | CN vs. MCI | MCI vs. LOAD | CN vs. LOAD | CN vs. LOAD |
| logFC | 0.207875239 | -0.180168831 | 0.20013891 | 0.068420156 | -0.020260737 | 0.282432909 | 0.22080636 | 0.16972131 |
| p-value | 1.23E-01 | 9.53E-01 | 5.13E-01 | 9.25E-01 | 1.00E+00 | 2.89E-01 | 3.26E-01 | 9.46E-02 |

**Table 2.**
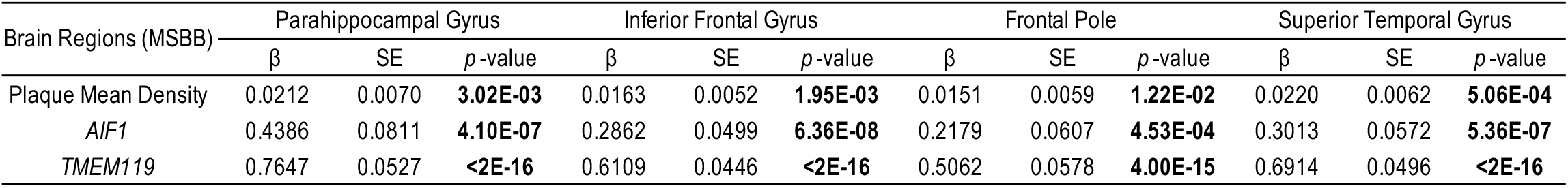
*INPP5D* expression levels are associated with amyloid plaque density and microglia-specific markers. Table 2 shows the β coefficient (β), standard error (SE), and *p*-value for the association analysis between *INPP5D* expression levels and amyloid plaque density or expression levels of microglia-specific markers *AIF1* and *TMEM119* by general linear models.

| Brain Regions (MSBB) | Parahippocampal Gyrus |  |  | Inferior Frontal Gyrus |  |  | Frontal Pole |  |  | Superior Temporal Gyrus |  |  |
| --- | --- | --- | --- | --- | --- | --- | --- | --- | --- | --- | --- | --- |
| | $\beta$ | SE | *p*-value | $\beta$ | SE | *p*-value | $\beta$ | SE | *p*-value | $\beta$ | SE | *p*-value |
| Plaque Mean Density | 0.0212 | 0.0070 | **3.02E-03** | 0.0163 | 0.0052 | **1.95E-03** | 0.0151 | 0.0059 | **1.22E-02** | 0.0220 | 0.0062 | **5.06E-04** |
| *AIF1* | 0.4386 | 0.0811 | **4.10E-07** | 0.2862 | 0.0499 | **6.36E-08** | 0.2179 | 0.0607 | **4.53E-04** | 0.3013 | 0.0572 | **5.36E-07** |
| *TMEM119* | 0.7647 | 0.0527 | **<2E-16** | 0.6109 | 0.0446 | **<2E-16** | 0.5062 | 0.0578 | **4.00E-15** | 0.6914 | 0.0496 | **<2E-16** |

### *INPP5D* expression levels are associated with amyloid plaque density in the human brain

We investigated the association between *INPP5D* expression levels and mean amyloid plaque densities in four brain regions (**Table 2**). Expression levels of *INPP5D* were associated with amyloid plaques in the parahippocampal gyrus (β=0.0212, p=3.02E-03; **Fig. 2a**), inferior frontal gyrus (β=0.0163, p=1.95E-03; **Fig. 2b**), frontal pole (β=0.0151, p=1.22E-02; **Fig. 2c**), and superior temporal gyrus (β=0.0220, p=5.05E-04; **Fig. 2d**).

**Fig. 2.**
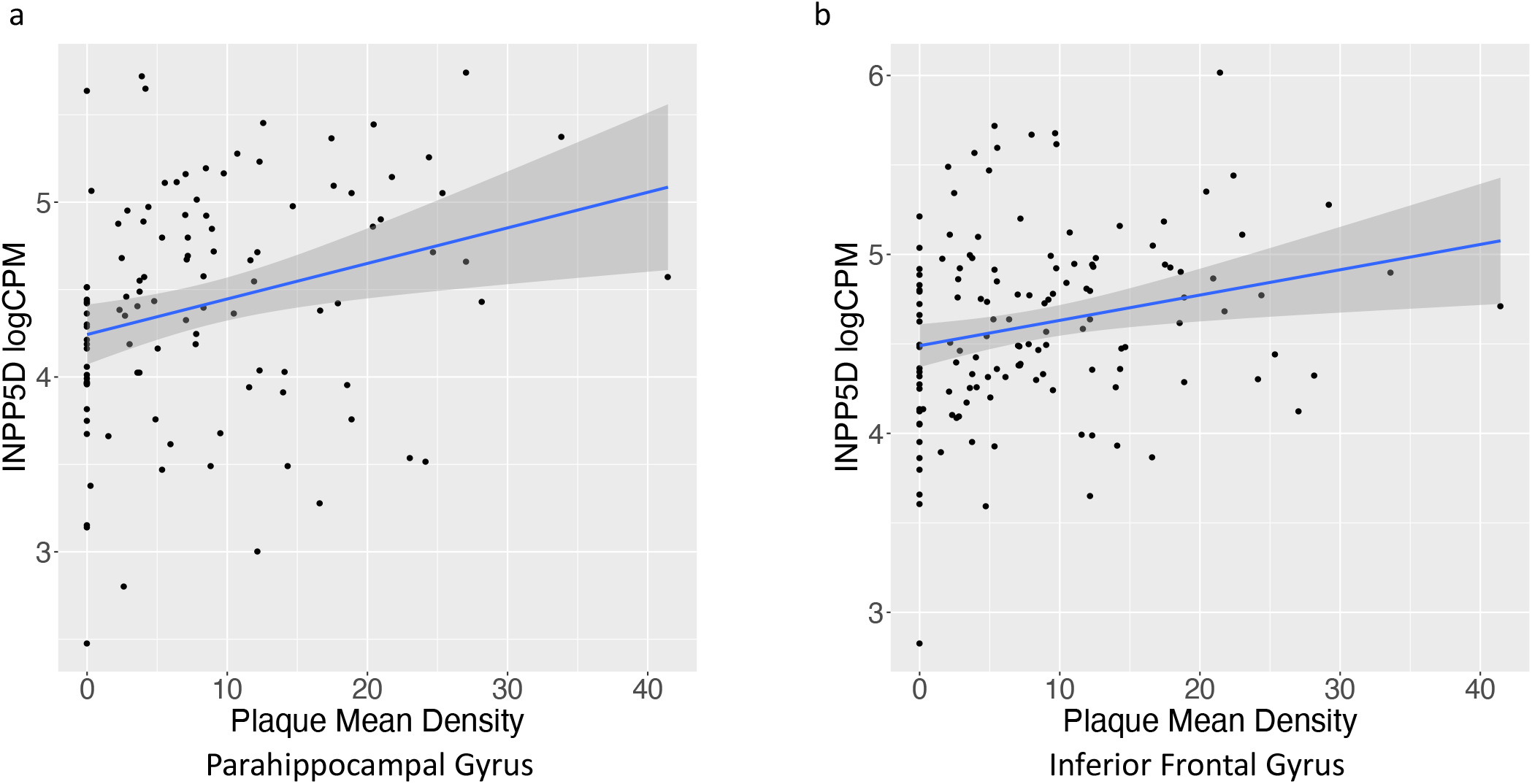
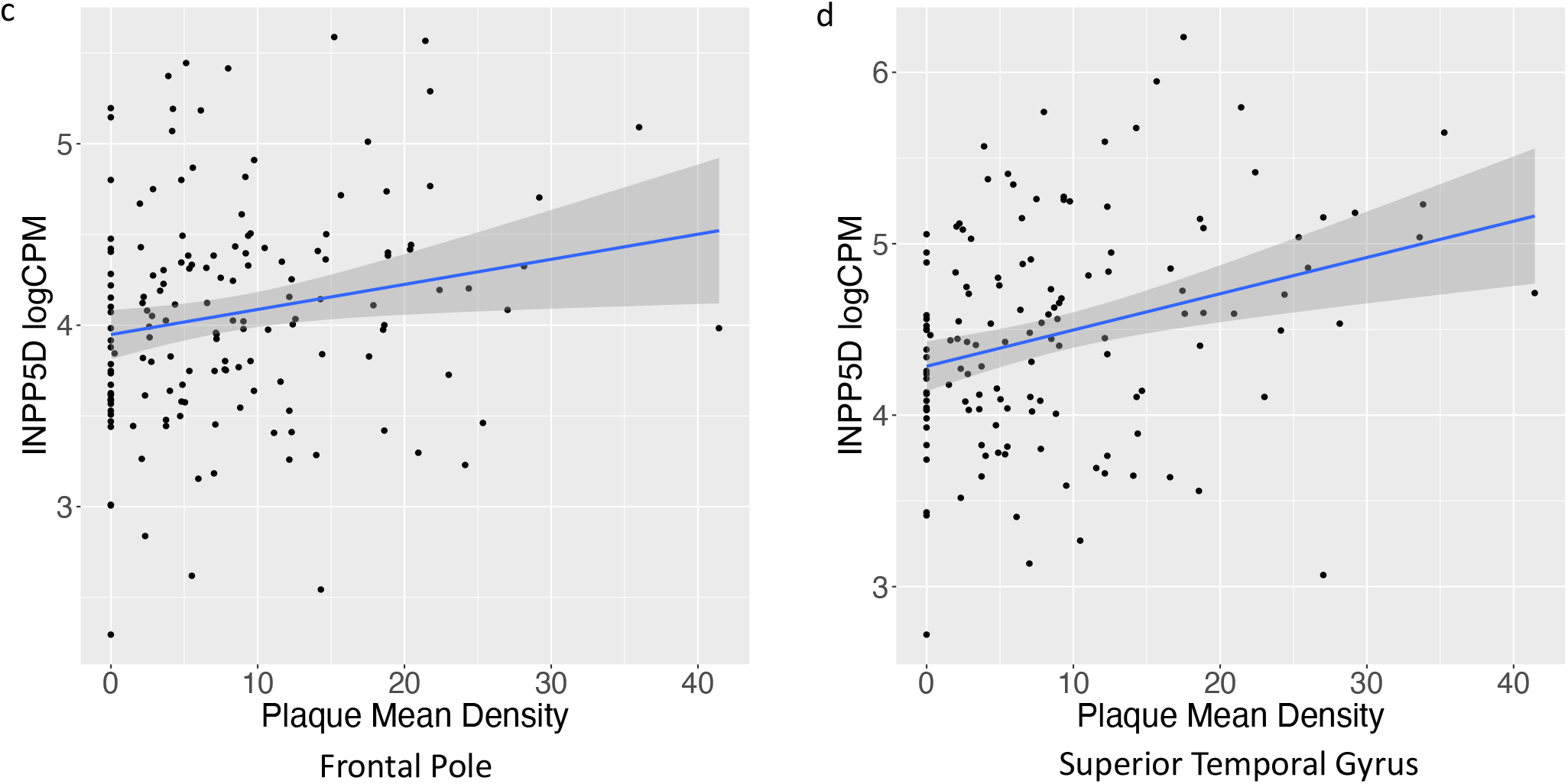
Association of *INPP5D* expression with amyloid plaque mean density. The scatter plots show the positive association between *INPP5D* expression and plaque mean density in (a) parahippocampal gyrus, (b) inferior frontal gyrus, (c) frontal pole, and (d) superior temporal gyrus from the MSBB cohort.

### *INPP5D* expression levels are increased in an amyloid pathology mouse model

We recapitulated our findings from the human data in the amyloidogenic mouse model, 5xFAD. We observed increased *Inpp5d* mRNA levels in 5xFAD mice throughout disease progression compared with WT controls in the brain cortex (**Fig. 3a**) and hippocampus (**Fig. 3b**) of four-, six-, eight-, and twelve-month-old mice (4-months: 1.57-fold in the cortex, 1.40-fold in the hippocampus; 6-months: 1.86-fold in the cortex, 2.61-fold in the hippocampus; 8-months: 2.23-fold in the cortex and 2.53-fold in the hippocampus; and 12-months: 1.93-fold in the cortex and 2.16-fold in the hippocampus). Similarly, INPP5D protein levels were increased in the cortex of 5xFAD mice at four and eight months of age (1.79 and 3.31-fold, respectively; p=0.06) (**Fig. 3c and 3d**). To assess *Inpp5d* induction was dependent on microglia, we depleted microglia in four-month-old 5xFAD mice by treating the animals with the colony-stimulating factor receptor-1 antagonist PLX5622 (PLX) for 28 days [18]. PLX treatment completely abolished the increase of *Inpp5d* in 5xFAD mice (**Fig. 3e**). Furthermore, expression levels of *Inpp5d* were restored after switching from the PLX diet to a normal diet for 28 further days (**Fig. 3f**).

**Fig. 3.**
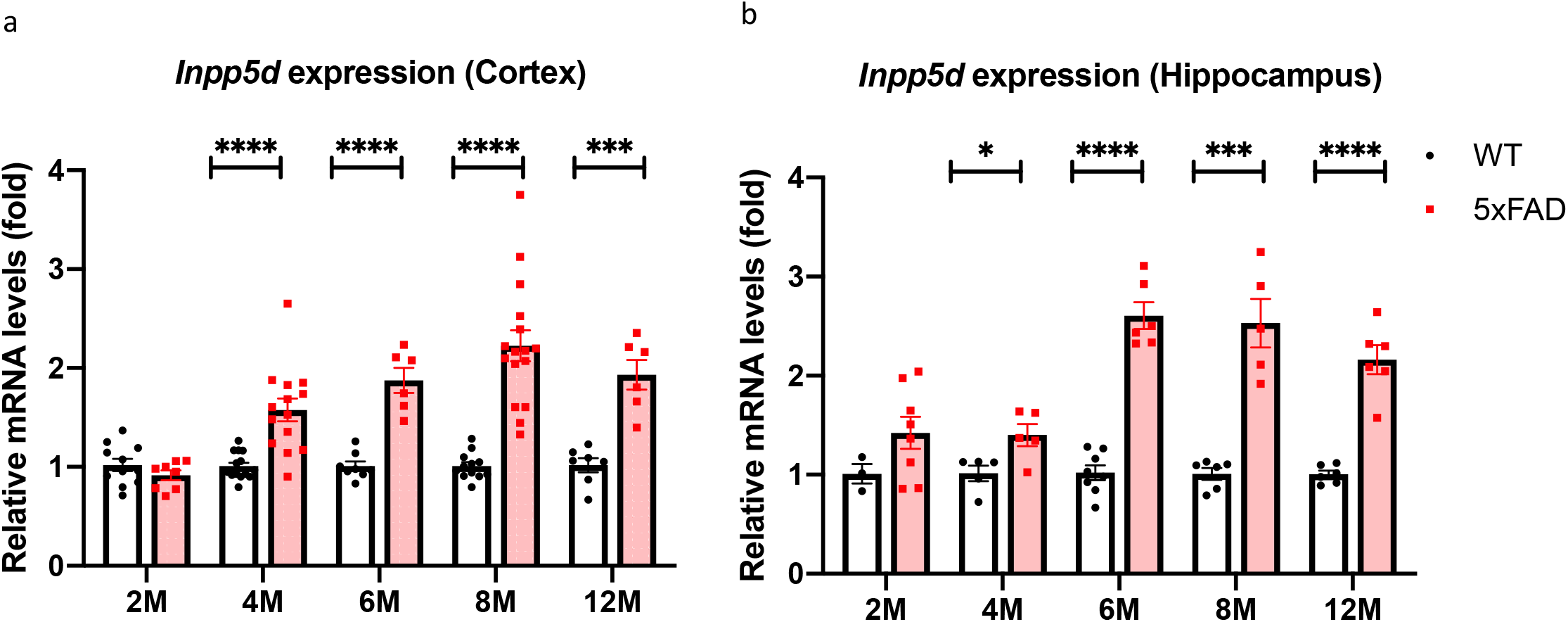
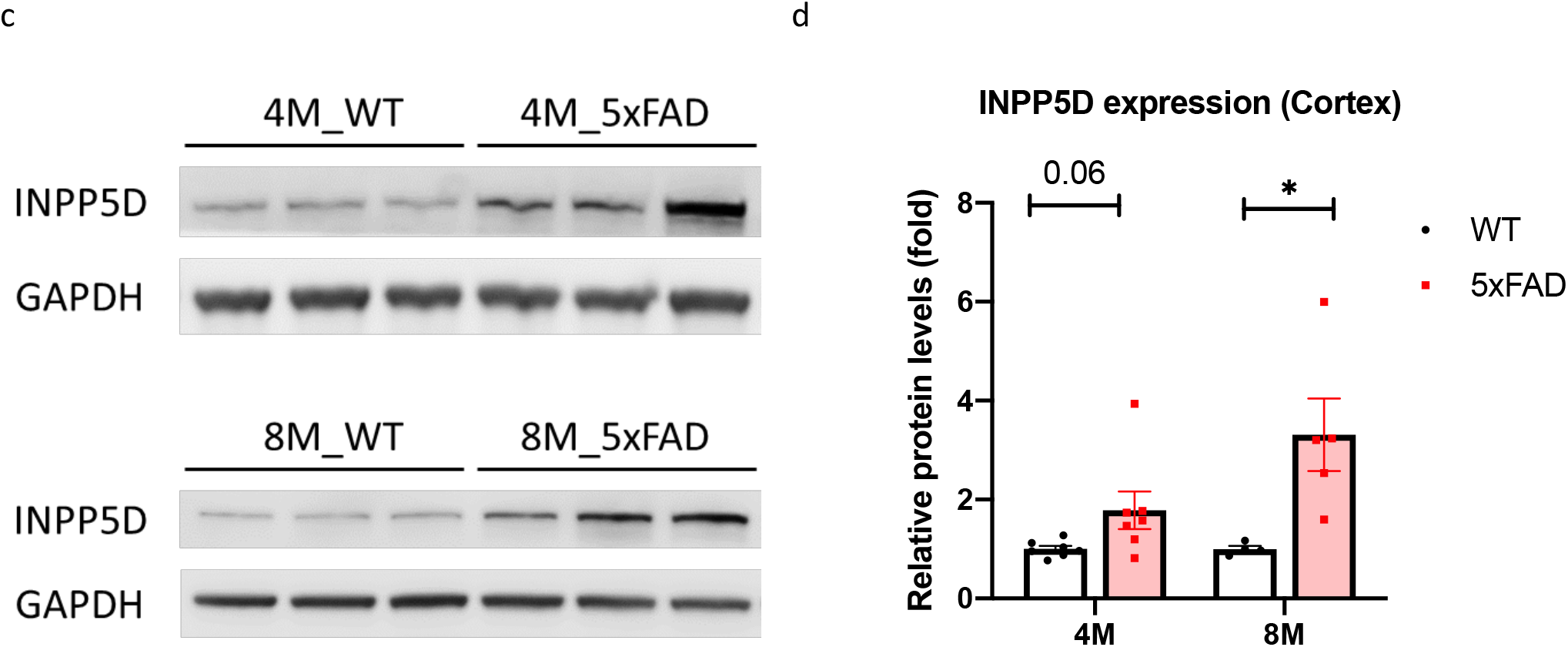
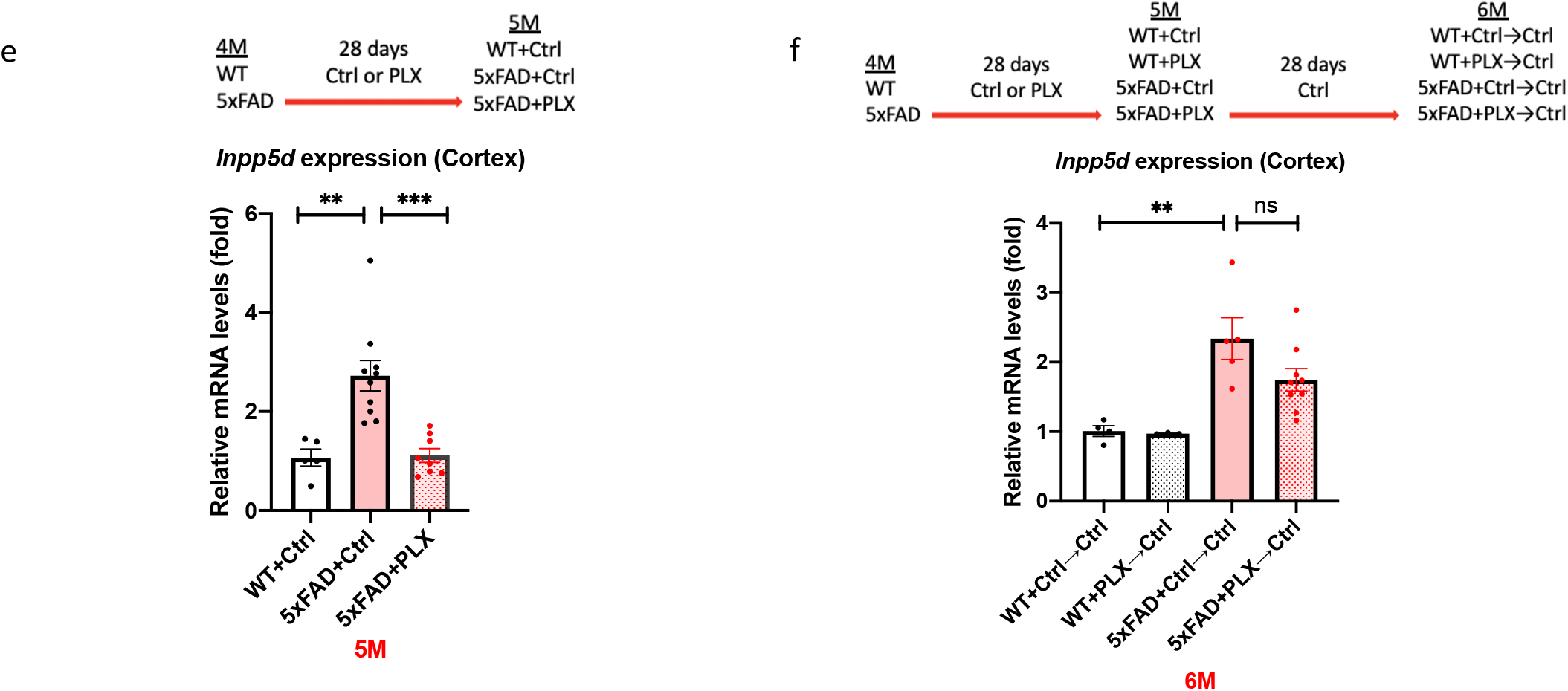
*Inpp5d* levels are increased in 5xFAD mice. Gene and protein levels of *Inpp5d* were assessed in cortical and hippocampal lysates from 5xFAD mice. Gene expression levels of *Inpp5d* were significantly increased in both cortex (a) and hippocampus (b) at 4, 6, 8, and 12 months of age (n=6-15 mice). There were significant changes in *Inpp5d* protein levels in the cortex at 8 months of age and an increased trend in the cortex at 4 months of age (n=4-7). Increased *Inpp5d* levels were abolished with PLX5622 treatment (e), and restored after switching PLX diet to normal diet (f) (n=3-10). *p<0.05; **p<0.01; ***p<0.001; ****p<0.0001, ns *not significant.*

### *INPP5D* expression levels are increased in plaque-associated microglia

Immunohistochemistry of 5xFAD mice brain slices at eight months old revealed that *Inpp5d* was mainly expressed in plaque-associated microglia (**Fig. 4**). INPP5D- and IBA1 (AIF1)-positive microglia cluster around 6E10-positive or X-34-positive plaques in the cortex (**Fig. 4a**) and subiculum (**Fig. 4b**). We did not detect any INPP5D expression in WT control mice (data not shown). Furthermore, analysis of transcriptomic data of sorted microglia from WT mouse cortex-injected labeled apoptotic neurons [12] revealed a reduction of *Inpp5d* expression levels in phagocytic microglia compared with non-phagocytic microglia (**Fig. 4c**), which is in agreement with the report that INPP5D inhibition promotes microglial phagocytosis [13].

**Fig. 4.**
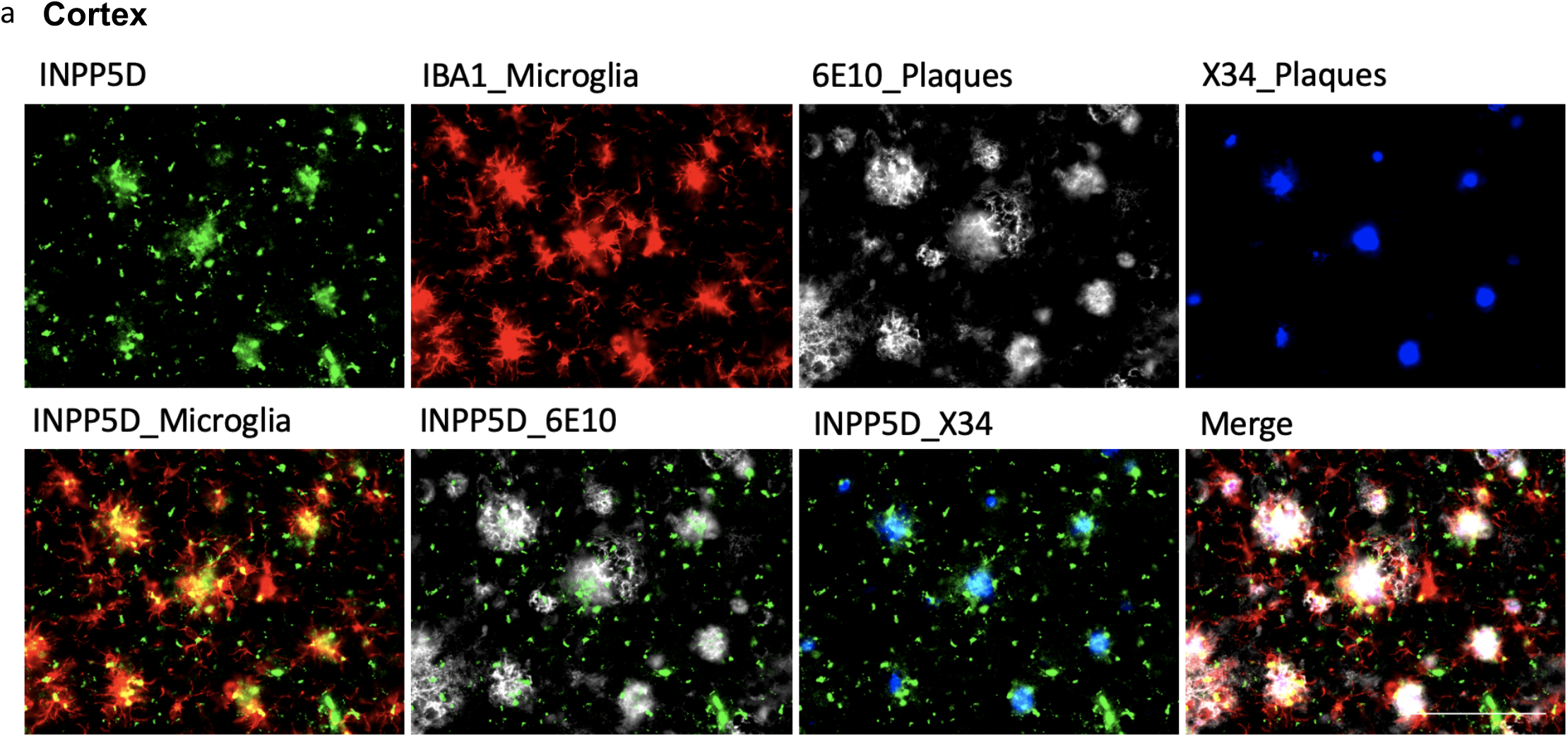
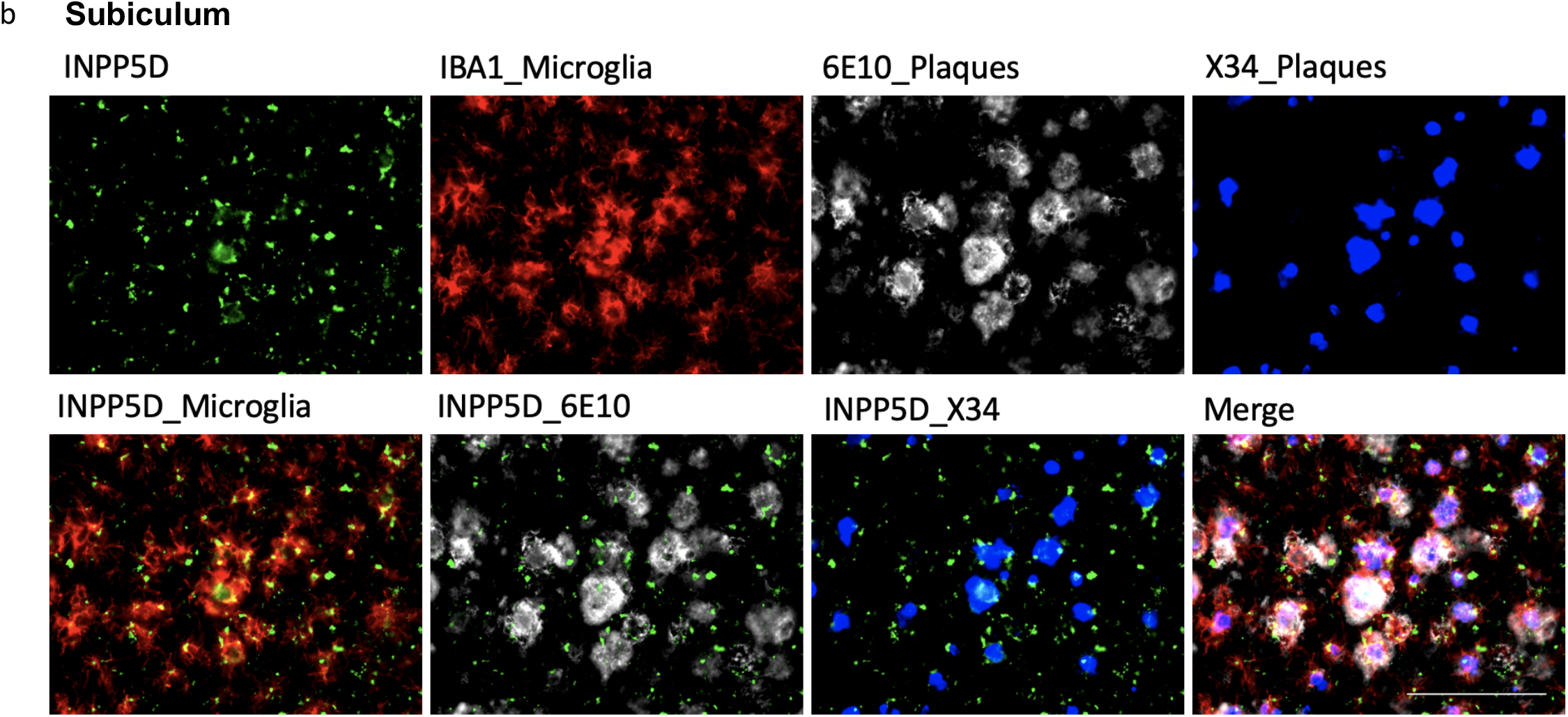
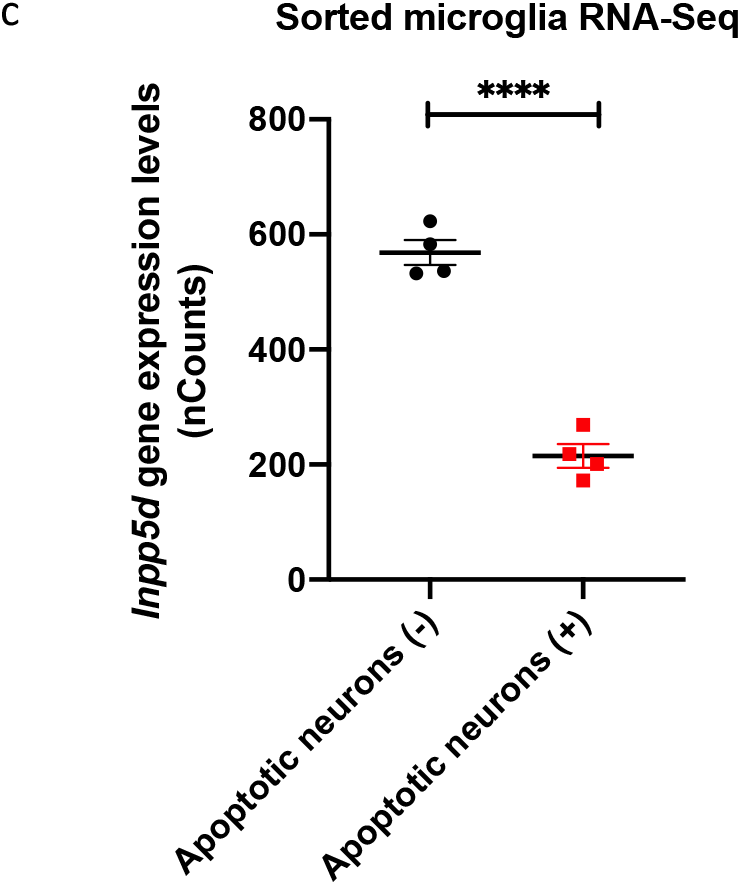
INPP5D expression levels were increased in plaque-associated microglia. INPP5D was mainly expressed in plaque-associated microglia. INPP5D- and IBA1 (AIF1)-positive microglia cluster around 6E10-positive or X-34-positive plaques in both cortex (a) and subiculum (b) of 8-month-old mice. Analysis of transcriptomic data of sorted microglia from wild-type mice cortex-injected labeled apoptotic neurons revealed that *Inpp5d* expression is increased in non-phagocytic microglia (Krasemann et.al) (c). Scale bar, 10 μm. ****p<0.0001

## Discussion

Although genetic variants in *INPP5D* have been associated with LOAD risk [5, 6, 20, 21], the role of *INPP5D* in AD remains unclear. We identified that *INPP5D* expression levels are increased in the brain of LOAD patients. Furthermore, expression levels of *INPP5D* positively correlate with brain amyloid plaque density and *AIF1* and *TMEM119* (microglial marker gene) expression [22–24]. We observed similar findings in the 5xFAD amyloidogenic model, which exhibited an increase in gene and protein expression levels of *Inpp5d* with disease progression, predominately in plaque-associated microglia, suggesting induction of *Inpp5d* in plaque-proximal microglia. Similarly, a recent study reported that *Inpp5d* is strongly correlated with amyloid plaque deposition in the APPPS1 mouse model [25, 26]. These findings are consistent with the observation of microgliosis in both AD and its mouse models.

INPP5D inhibition has been associated with microglial activation and increased phagocytic activity, which is consistent with our transcriptomic data of sorted microglia from murine brains injected with apoptotic neurons [12], showing a decrease in *Inpp5d* expression in phagocytic microglia compared to non-phagocytic. These findings support the hypothesis that an increase in *INPP5D* expression in AD is a part of an endogenous homeostatic microglial response to negatively control their own activity. However, this “brake” might be excessive in AD, as reflected in our findings that *INPP5D* expression is elevated in LOAD. *INPP5D* overexpression might result in microglia with deficient phagocytic capacity, resulting in increased Aβ deposition and neurodegeneration. Thus, the pharmacological targeting of *INPP5D* might be a novel therapeutic strategy to shift microglia towards a beneficial phenotype in AD. Future studies in genetic mouse models are necessary to further clarify the role of *INPP5D* in microglial function and AD progression.

## Conclusions

In conclusion, our results demonstrate that *INPP5D* plays a crucial role in AD pathophysiology and is a potential therapeutic target. *INPP5D* expression is upregulated in LOAD and positively correlated with amyloid plaque density. *Inpp5d* expression increases in the microglia of 5xFAD mice as AD progresses, predominately in plaque-associated microglia. Future studies investigating the effect of *INPP5D* loss-of-function on microglial phenotypes and AD progression may allow for the development of microglial-targeted AD therapies.

## List of abbreviations

AD: Alzheimer’s disease
LOAD: late-onset AD
GWAS: genome-wide association studies
INPP5D: phosphatidylinositol 3,4,5-trisphosphate 5-phosphatase 1
PI(3,4,5)P3: phosphatidylinositol (3,4,5)-trisphosphate
PI(3,4)P2: phosphatidylinositol (3,4)-bisphosphate
CSF: cerebrospinal fluid
OR: odds ratio
CI: confidence interval
β: β coefficient
WT: wild-type
MCI: mild cognitive impairment
APOE ε4: apolipoprotein ε4 allele
PFA: paraformaldehyde
PCR: polymerase chain reaction
Seq: sequencing
ANOVA: analysis of variance
qPCR: quantitative real-time PCR
mAb: monoclonal antibody.

## Declarations

### Ethics approval and consent to participate

Animals used in the study were housed in the Stark Neurosciences Research Institute Laboratory Animal Resource Center at Indiana University School of Medicine and all experimental procedures were approved by the Institutional Animal Care and Use Committee.

### Consent for publication

All participants were properly consented for this study.

### Availability of data and materials

The datasets analyzed during the current study are available from the corresponding author on reasonable request.

### Competing interests

The authors declare that they have no competing interests.

### Funding

This work was supported by NIA grant RF1 AG051495 (B.T.L and G.E.L), NIA grant RF1 AG050597 (G.E.L), NIA grant U54 AG054345 (B.T.L et al.), NIA grant K01 AG054753 (A.L.O), NIA grant R03 AG063250 (K.N), and NIH grant NLM R01 LM012535 (K.N)

### Author contributions

A.P.T, P.B.L, C.D, Y.L, B.T.L, G.E.L, A.L.O, and K.N designed the study. A.P.T, P.B.L, C.D, M.M, B.T.C, and K.N performed the experiments and analyzed the data. A.P.T, M.M, G.E.L, A.L.O, and K.N wrote the manuscript. All authors discussed the results and commented on the manuscript.

## Acknowledgments

We would like to thank Dr. Richard W. Cho at Cell Signaling Technology for providing SHIP1/INPP5D Rabbit mAb. We thank Louise Pay for critical comments on the manuscript. We thank Cynthia M. Ingraham, Deborah D. Baker, Christopher D. Lloyd, Stephanie J. Bissel, Shweta S. Puntambekar, Guixiang Xu, Roxanne Y. Williams, and Teaya N. Thomas for their help with taking care of the mice, genotyping, and helpful discussions. We thank the International Genomics of Alzheimer’s Project (IGAP) for providing summary results data for these analyses. IGAP investigators contributed to the design and implementation of IGAP and/or provided data but did not participate in analysis or writing of this report. IGAP was made possible by the generous participation of the control subjects, the patients, and their families. The i–Select chips were funded by the French National Foundation on Alzheimer’s disease and related disorders. EADI was supported by the LABEX (laboratory of excellence program investment for the future) DISTALZ grant, Inserm, Institut Pasteur de Lille, Université de Lille 2, and the Lille University Hospital. GERAD/PERADES was supported by the Medical Research Council (Grant n° 503480), Alzheimer’s Research UK (Grant n° 503176), the Wellcome Trust (Grant n° 082604/2/07/Z) and German Federal Ministry of Education and Research (BMBF): Competence Network Dementia (CND) grant n° 01GI0102, 01GI0711,01GI0420. CHARGE was partly supported by the NIH/NIA grant R01 AG033193 and the NIA AG081220 and AGES contract N01–AG–12100, the NHLBI grant R01 HL105756, the Icelandic Heart Association, and the Erasmus Medical Center and Erasmus University. ADGC was supported by the NIH/NIA grants: U01 AG032984, U24 AG021886, U01 AG016976, and the Alzheimer’s Association grant ADGC–10–196728.

## Notes

### Competing Interest Statement

The authors have declared no competing interest.

